# Altered topology of neural circuits in congenital prosopagnosia

**DOI:** 10.1101/100479

**Authors:** Gideon Rosenthal, Michal Tanzer, Erez Simony, Uri Hasson, Marlene Behrmann, Galia Avidan

## Abstract

Using a novel fMRI-based inter-subject functional correlation (ISFC) approach, which isolates stimulus-locked inter-regional correlation patterns, we compared the cortical topology of the neural circuit for face processing in participants with congenital prosopagnosia (CP) and matched controls. Whereas the anterior temporal lobe served as the major network hub for face processing in controls, this was not the case for the CPs. Instead, this group evinced hyper-connectivity in posterior regions of the visual cortex, mostly associated with the lateral occipital and the inferior temporal cortices. Moreover, the extent to which the network organization was atypical differed as a function of the severity of the face recognition deficit. These results offer new insights into the perturbed cortical topology in CP, which may serve as the underlying neural basis of the behavioral deficits typical of this disorder. The approach adopted here has the potential to uncover altered topologies in other neurodevelopmental disorders, as well.

**Significance Statement:** Congenital prosopagnosia (CP; ‘face blindness’), a developmental deficit in face recognition, is thought to affect up to 3% of the population. Understanding its neural basis is challenging as there is no obvious deficit on conventional structural or functional MRI scans. Using an innovative, fMRI-based inter-subject correlation approach geared towards tracking inter-regional stimulus-locked brain activation, the present study uncovers marked topological differences in a distributed brain network of higher-order visual regions in CP relative to controls. Alteration in topology also differs as a function of the severity of the deficit. These findings shed new light on the neural perturbations underlying CP, and the analytic approach we have adopted may have utility in elucidating the neural basis of other neurodevelopmental disorders such as dyslexia or amusia.

## Introduction

Understanding the neural basis of developmental disorders such as congenital prosopagnosia (CP) remains a challenge from both a basic science and a translational perspective as there are no obvious identifiable deficits on conventional anatomical MR brain scans. Furthermore, many studies show that CP individuals evince normal fMRI activation in the ‘core’ face-related posterior patches of the brain (Hasson et al., 2003; Avidan et al., 2005, 2014; Avidan and Behrmann, 2009) (but see von Kriegstein et al., 2008; Dinkelacker et al., 2011; Furl et al., 2011). In contrast, more sensitive methods that have been used to map structural changes in CP relative to controls, such as diffusion tensor imaging (DTI) have revealed a reduction in long-range white matter tracts connecting the ‘core’ face-related posterior patches and the anterior temporal lobe face patch (ATL) in CP (Behrmann et al., 2007; Steinbrink et al., 2008; Odegard et al., 2009; Thomas et al., 2009). Other studies have also reported local structural and functional atypical alterations in the vicinity of face-selective regions (Gomez et al., 2015; Song et al., 2015; Lohse et al., 2016). Using standard functional connectivity (FC) analysis, which measures the temporal correlations across different brain areas within an individual, we have previously documented impairments in the connectivity patterns between the ‘core’ and ‘extended’ nodes of the face system (Avidan and Behrmann, 2009, 2014; Avidan et al., 2014).

The pattern of FC within each individual, as utilized in previous studies, is a combination of stimulus-induced correlations, intrinsic neural fluctuations, and correlations induced by non-neuronal artifacts (such as head motion, respiration). Separating these factors is challenging within the framework of standard FC, given the strength of the intrinsic neural fluctuations. Hence, group differences in FC may not be sufficiently robust to be detected following whole brain statistical correction, and, therefore, are less suitable for mapping large-scale changes in network topology.

In contrast with these previous studies that have examined the neural profile of CP based on a subset of brain regions and their connectivity, here, we adopt an innovative, large-scale network approach. The primary goal of this approach is to elucidate the functional brain topology in individuals with CP (and matched controls) as a means of examining alterations in neural circuitry. Because there is consensus that multiple regions (face ‘patches’) are implicated in normal face recognition in humans (Haxby et al., 2000; Pyles et al., 2013; Weiner and Grill-Spector, 2013) and in non-human primates (Tsao et al., 2006; Hung et al., 2015), elucidating alterations in the topology of this distributed cortical circuit is of great interest.

To that end, in the present study, we have used a novel method, termed “inter-subject functional correlation” (ISFC), which is designed to isolate stimulus-locked functional responses, by correlating the response profile across the brains of multiple participants (Simony et al 2016). Importantly, intrinsic neural dynamics during rest and task conditions that are not related to the pattern of activation evoked by stimulation, as well as non-neuronal artifacts (e.g., respiratory rate, motion), only influence the pattern of correlations within each individual brain, but cannot induce correlations between subjects. In contrast, neural processes that are locked to the structure of the stimulus can be correlated across brains. Thus, the ISFC method allows us to track stimulus-locked brain responses within the high-level visual network in control subjects and to contrast these correlation patterns with the patterns uncovered in CP individuals. Such an approach allows us to explore possible alterations in connectivity across large swaths of the cortex in an assumption-free manner rather than focusing on a predetermined subset of regions and connections.

## Results

During an fMRI scan, 10 CP and 10 control subjects viewed separate blocks of images of emotional faces (angry, fearful), neutral faces, famous faces, unfamiliar faces and buildings (Avidan et al., 2014). To define an initial set of unbiased nodes (functional regions or clusters comprising the network) with which to explore topology and connectivity, face-selective (right FFA) and non-face selective (right LOC) seed regions were defined based on BOLD data from a separate group of 16 control subjects (see Methods). Using these seed regions, within each individual, two correlation maps were derived and a binary mask was constructed for each. These masks were then separately sub-divided into small, spatially constrained clusters with each cluster serving as a node for the network analyses. This procedure resulted in large swaths of cortex sub-divided into nodes, each of which preserved the original "functional tagging" from the seed correlation analysis (i.e., face-selective voxels, orange color, object-selective voxels, green color, or voxels that were common to both maps, blue color; see Methods for details on node definition, see Figure 1).

**Figure 1.**
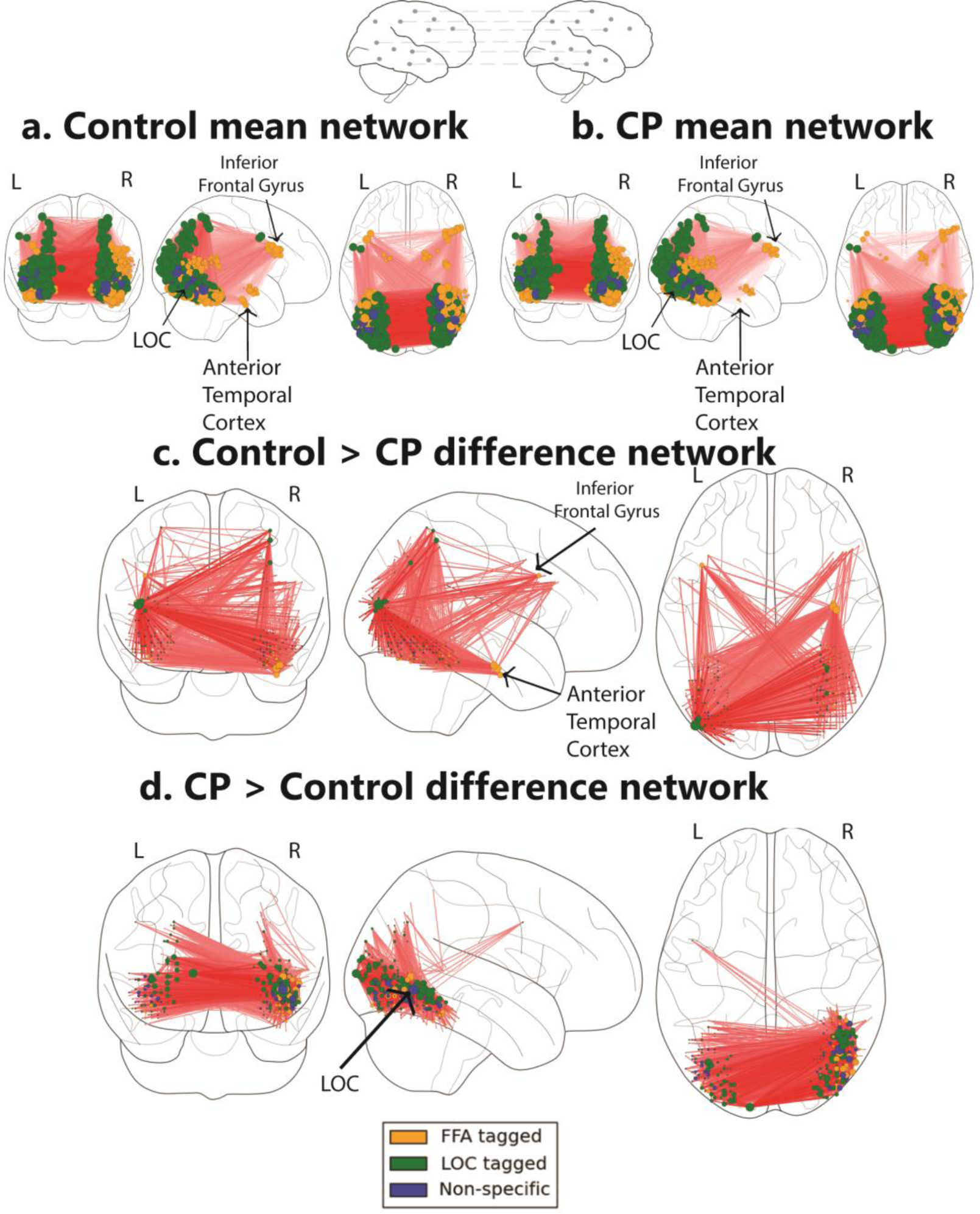
Networks obtained using the ISFC approach. **a.** Mean network obtained from the control group **b.** Mean network obtained from the CP group. These mean networks are shown for visualization purposes. For each group, networks are projected on three views of the brain (left and right lateral views and a ventral view). The colors of the nodes reflect their functional selectivity (face-selective, non-face selective, and nodes which are not selective to either of these stimuli). The size of the nodes is proportional to their degree (the larger the node, the greater its ISFC). The color of the edges reflects the weight of the connections such that darker connections have higher values. The same conventions are used in all figures. **c.** The difference network of controls>CPs. This comparison is presented using 0.3 edge threshold, which reflects the difference between the correlation coefficient values of the controls compared to the CP individuals. The ATL and the inferior frontal gyrus are marked. As is graphically depicted, the ATL serves as the main hub across thresholds for controls but not for CP. Three nodes, which comprise the ATL, were ranked in the top 10 ranks of degree scores (1-3; see Table 3a). **d.** Difference networks obtained from the comparison of CPs > controls. CPs evince hyper-ISFC in posterior visual regions. This is evident by the elimination of edges as the threshold increases and by the color spectrum of the anterior vs. posterior edges reflecting their weight.

**Table 3.** *Rank of top 10 nodes obtained from the comparisons of CP and the control group:*

| **a. Controls>CPs** |  | **b. CPs>Controls** |  |
| --- | --- | --- | --- |
| *Rank* | *Region Name* | *Rank* | *Region Name* |
| 1,2,3 | Anterior temporal Cortex (faces) | 1,3,7,8,9,10 | Right inferior temporal gyrus (non-faces) |
| 4,8 | Left inferior frontal gyrus opercular part (faces) | 2 | Left lateral occipital cortex (non-faces) |
| 5 | Left TOS (non-faces) | 4,6 | Right lateral occipital cortex (non-faces) |
| 6 | Right FFA (faces) | 5 | Right lateral occipital cortex (faces) |
| 7,10 | Right inferior frontal gyrus opercular part (faces) |  |  |
| 9 | Amygdala left (faces) |  |  |

The 10 nodes with the highest node degree obtained from the Controls>CPs (**a.**) and CPs>Controls (**b.**) difference networks. Anatomical locations, which are based on "the atlas of human brain" (Mai et al., 2008) and validated by an expert are provided for each node. Note that in the controls>CP network most of the nodes were face selective, while in the CP>controls network, only a single node was face selective.

### Inter-subject functional correlation *(ISFC) analysis*

To detect stimulus-locked changes in functional responses to faces in CP relative to controls, we used the inter-subject functional correlation (ISFC; see Methods), which calculates the inter-regional correlations in the brains of different individuals who viewed the same stimuli (Simony et al 2016). Simony et al. (2016) demonstrated that the ISFC method substantially increases the signal-to-noise (SNR) ratio in detecting shared stimulus-locked network correlation patterns by filtering out idiosyncratic intrinsic neural dynamics and non-neuronal artifacts (e.g., respiratory rate; motion) that can influence FC patterns within a brain but that are uncorrelated across the brains of different participants. Capitalizing on the high SNR of the ISFC procedure permits the construction of a fine-grained functional brain network even with a relatively small sample size as is the case in the present study and potentially in other situations of relatively rare disorders (see Figure 5 and Methods for ISFC workflow).

We calculated the ISFC within the CP and the control groups using BOLD data activated in response to faces and buildings (see below). For comparison, we also ran the same analysis using a standard FC procedure. In the FC analysis, the response profile in each ROI was correlated with the response profile of all other ROIs within an individual. The analysis was repeated for each individual in each group and statistical significance for each edge was determined using t-test followed by FDR correction. We return to these results below. ISFC is similar in logic to FC, with one critical difference: instead of correlating the response profile within the brain of each individual, we calculated the correlation patterns across brains. Each experimental group (controls and CPs) was randomly split into 2 halves 1000 times (see split-group ISFC procedure in Materials and Methods) and the average non-thresholded networks for each group are presented in Figure 1a,b for visualization purposes.

As is evident, the raw mean networks over the split-group ISFC of each group are visually similar. To evaluate the statistical differences of the network structure across the two groups, the networks were directly compared using a permutation test (see Materials and Methods for details). This resulted in a difference network in which the edges indicate the significant difference between the two groups (either controls>CPs or CPs>controls). Non-significant edges were eliminated (see details of statistical analysis in Methods ISFC Formulation section and Figure 5).

The control>CP difference network revealed that control participants, but not CP, exhibited increased ISFC patterns from nodes in the vicinity of the ATL both to face and non-face selective nodes located throughout the visual cortex. This effect was substantial and apparent at all thresholds (see specific pattern of ATL ISFC in Table 2). (Figure 1 c, d). The group differences were further quantified using a measure of ‘node degree’, which quantifies the number of edges connecting to a node (Rubinov and Sporns, 2010). The degree scores of all nodes in the network were ranked in a descending order, and, of interest, the three nodes located in the ATL were ranked as the top 3 for the control subjects (see Figure 1c,d and Table 3a). Note that the ATL served as a hub connecting both face and non-face selective nodes (see Table 2). The node rankings confirm the centrality of the ATL in the face network of controls in contrast with that of CP, whose top 3 rankings include the right inferior temporal gyrus and the left lateral occipital cortex.

**Table 2.** *Coordinates of the ATL nodes, and the calculated values of the within module degree and participation coefficient*

| Number<br>Of Voxels | MNI<br>center<br>coordinates | Within<br>module<br>degree | Participation<br>coefficient |
| --- | --- | --- | --- |
| 23 | 42, -14, -26 | 5.03 | 0.62 |
| 13 | 40, -10, -32 | 4.40 | 0.60 |
| 20 | 40, -12, -28 | 5.62 | 0.62 |
| **<u>Voxels</u>** | 40, -10, -32 | **<u>Weighted</u>** | **<u>Weighted</u>** |
| **<u>Sum:</u>** |  | **<u>Mean:</u>** | **<u>Mean:</u>** |
| 55 |  | 5.09 | 0.61 |

The network obtained from the CPs>controls comparison revealed that, at a low-edge threshold, significant edges were located throughout the visual cortex, but, as the threshold was increased, edges from anterior regions were eliminated and the remaining significant edges were located only in posterior parts of the visual cortex (Figure 2 b). To further quantify this effect, the Y coordinates of the 3D MNI space were ranked in an ascending order and binned into 21 equally sized bins measuring distance parallel to the posterior-anterior commissure axis (the number of bins was chosen to be maximal with a constraint that each bin contains at least one node). The significance level of the posterior to anterior pattern was then quantified using the Spearman correlation between the Y coordinate bins and the nodes’ degree. The degree value, at an edge difference threshold of 0, was then calculated for these bins. The results indicate that the lower Y coordinates (more posterior regions along the anterior posterior axis) were associated with higher degree values (r_s_ = −0.63, P < 0.005; Figure 2b), validating the greater posterior ISFC in CP versus controls. Examining the controls>CPs contrast revealed an opposite pattern, with greater anterior compared to posterior ISFC (r_s_ = 0.73, P < 0.005; Figure 2a).

**Figure 2.**
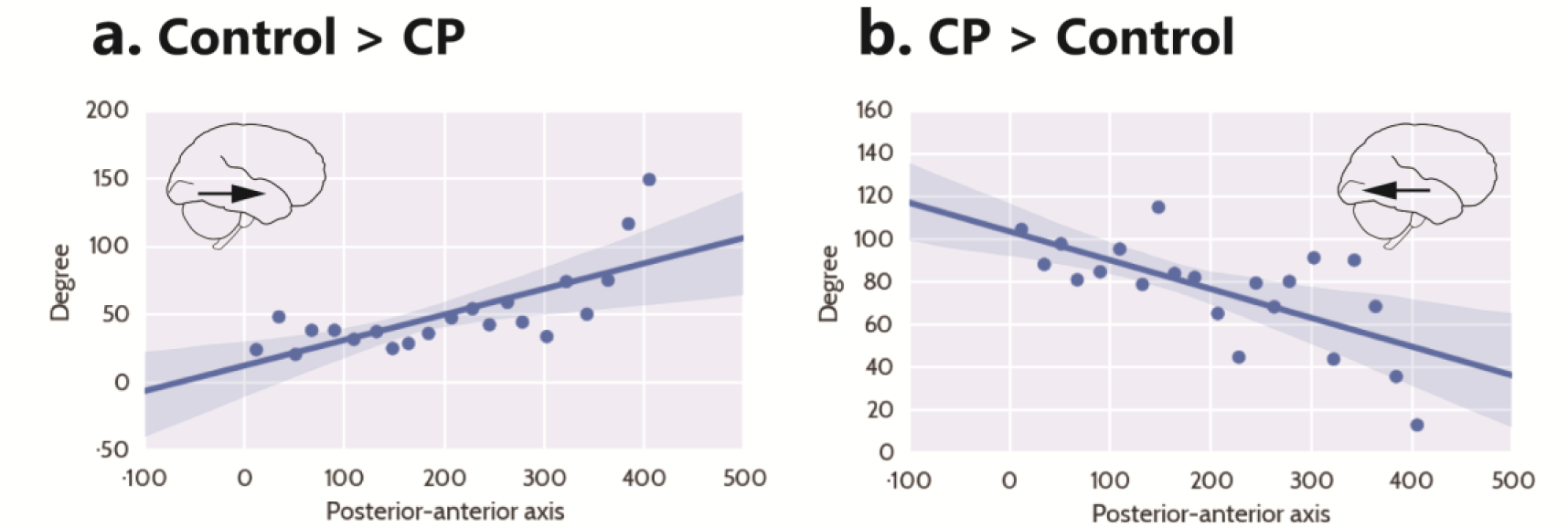
ISFC correlation between degree and node location along posterior-anterior axis. **a.** Linear regression fit with a 95% confidence interval band between the degree measure of the controls>CPs network nodes, at minimum threshold, and the Y coordinate ascending rank order of each node binned into 21 equally sized bins (r_s_ = 0.73, P < 0.005). The x-axis of the graph denotes the Y coordinates of the 3D MNI space ranked in ascending order, the y-axis of the graph marks the degree value of each node. As is evident, the higher the y-coordinate (more anterior), the higher the degree value. **b.** Linear regression fit with a 95% confidence interval band between the degree measure of the CPs>controls network nodes, at minimum threshold, and the Y coordinate ascending rank order of each node binned into 21 equally sized bins (r_s_ = −0.63, P < 0.005). As is evident, the relationship is negative: the higher the y-coordinate (more anterior), the lower the degree value.

Additionally, this analysis revealed face-specific dominance, such that the nodes that had the highest degree were face-selective. Specifically, higher ISFC patterns in the control group, compared to the CP group, were associated with face selective nodes. For this comparison, nine out of the top ten nodes (90%) were face-related (ranked in descending order by degree score) (Table 3a), compared to an overall face node base rate of 35% (number of face nodes divided by overall number of nodes). The difference between the face-selective node rate in the control>CP contrast and the overall face nodes base rate was statistically significant χ^2^(1, n=20)=13.2, p<.001. Thus, the face network of the controls was associated with a higher number of face-tagged nodes compared with the face network of the CPs. When comparing the opposite contrast (CP>controls), no statistically significant difference was found in the face-selectivity of the nodes χ^2^(1, n=20)=2.74, p=n.s. In fact, four of the top ten nodes, of which only one was face specific, belonged to the lateral occipital cortex (one in the left hemisphere and three in the right) and six nodes belonged to the adjacent right inferior temporal cortex.

Finally, given the known dominance of the right hemisphere in face processing (Haxby et al., 2000; Jonas et al., 2012; Parvizi et al., 2012; Rossion, 2014), we compared the network characterization across the two hemispheres by applying a Mann-Whitney test on the node degree measure obtained from the controls>CP and CP>controls contrasts. While no significant difference was found for the controls>CP contrast in face-tagged nodes (Median = 36, Median =45.5 in right vs. left hemisphere U = 3702, p = n.s), examining the CP>controls contrast revealed that the degree values were greater for the right (Median = 23.5) than left hemisphere (Median = 2) in face-tagged nodes, U = 2275, p = 0.0005, r =0.28. This indicates that, in the CP>controls, but not in the controls>CP contrast, greater ISFC was evident in the right versus left hemisphere in face-tagged nodes. This finding implies that the altered topological organization was more pronounced in the right than left hemisphere of the CPs in face-tagged nodes.

### Functional correlation analysis

We next compared the results obtained using ISFC to the results obtained using standard functional connectivity (FC) analysis. The difference in FC patterns between CP and the control subjects was small relative to the ISFC analysis. In accordance with our prediction, when comparing the controls to CPs, we observed the expected greater connectivity between the ATL and posterior regions. However, this effect was weak and was not evident following the application of the FDR multiple comparisons procedure (see Figure 3a, controls>CPs). Similar to the ISFC, the opposite contrast of CPs>controls revealed hyper connectivity in posterior visual regions with a main hub in the vicinity of the LOC and inferior temporal cortex, although this was weaker. Critically, this effect was also evident following FDR correction (see Figure 3b, CPs>controls).

**Figure 3.**
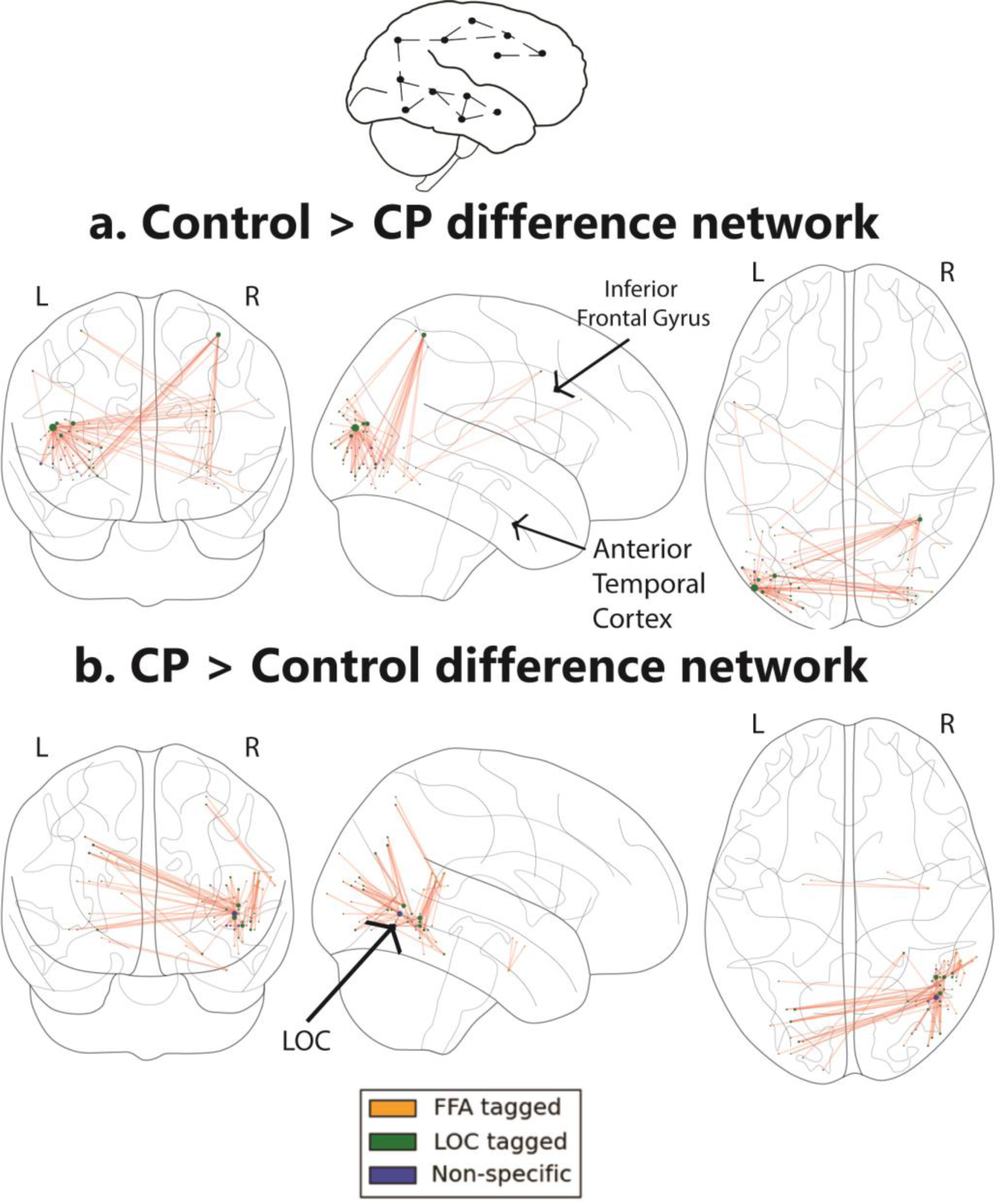
Networks obtained using the FC approach. **a.** The FC difference network of controls>CPs. This comparison reflects the difference between the FC correlation coefficient values of the controls compared to the group of CP individuals. The maps are presented following the application of the FDR correction (q<0.05). Note that the ATL related edges are not statistically significant after FDR correction. **b.** Difference networks obtained from the comparison of FC in CPs > controls.

#### Severity of face recognition deficits and network topology in CP

Thus far, the findings reveal that, relative to controls, the CPs exhibited atypical network topology with reduced ISFC to the ATL hub and increased ISFC more posteriorly, specifically to the LOC and inferior temporal cortex. To assess whether these topological changes are functionally related to the severity of the prosopagnosia, we split the CP group into more mildly and more severely affected sub-groups, based on the behavioral scores on the CFMT (see Table 1) and repeated the analyses used above. Compared to controls, both the severe and mild CP sub-groups had reduced ISFC patterns with the ATL at all thresholds (Table 5a and 6a). Unsurprisingly, the rankings of the ATL confirmed its centrality in the face network of the controls in contrast with each of the CP subgroups: also, comparisons of controls>severe CP and controls>mild CP contrast revealed greater anterior compared to posterior ISFC (r_s_ = 0.88, P < 0.005 and r_s_ = 0.85, P < 0.005 respectively). Contrasting each of the two CP groups against the controls indicates that more posterior regions along the anterior posterior axis are associated with higher degree values [CP severe>controls (r_s_ = −0.48, P < 0.05); CP mild>controls (r_s_ = −0.5, P < 0.05)]. Thus, we replicated the effect in both the severe and mild sub-groups and, similar to the comparison of the entire CP group to the controls, a posterior to anterior pattern emerged such that the more posterior a node, the higher its degree.

**Table 1.** *CP behavioral scores ordered by severity as indicated by performance on the CFMT*

|  | Participant | Sex | Age | Famous faces questionnaire |  | CFMT (total) |  |
| --- | --- | --- | --- | --- | --- | --- | --- |
|  |  |  |  | % corr. | Z- score |  |  |
| Severe CP | BL | F | 18 | 20 | -4.88 | 28 | -4.15 |
|  | BQ | F | 29 | 16 | -4.92 | 30 | -3.89 |
|  | KG | F | 49 | 75 | -0.69 | 33 | -3.5 |
|  | ON (Avidan et al., 2014) | F | 48 | 60.7 | -1.77 | 35 | -3.23 |
|  | MT<br>(Nishimura et al., 2010),<br>(Avidan et al., 2011),<br>(Behrmann and Avidan, 2005),<br>(Thomas et al., 2009),<br>(Avidan and Behrmann, 2008),<br>(Humphreys et al., 2007) ,<br>(Behrmann et al., 2007),<br>(Avidan et al., 2014) | M | 50 | 62.5 | -1.64 | 36 | -3.11 |
| Mild CP | WA<br>(Nishimura et al., 2010),<br>(Avidan et al., 2014) | F | 23 | 45.7 | -2.91 | 40 | -2.58 |
|  | KE (Avidan et al., 2011),<br>(Thomas et al., 2009),<br>(Avidan et al., 2014) | F | 67 | 42.9 | -3.12 | 40 | -2.58 |
|  | TD<br>(Nishimura et al., 2010),<br>(Avidan et al., 2011),<br>(Avidan et al., 2014) | F | 38 | 46.4 | -2.85 | 41 | -2.45 |
|  | MN<br>(Nishimura et al., 2010),<br>(Avidan et al., 2014) | F | 50 | 60.7 | -1.77 | 52 | -1 |
|  | BT (Avidan et al., 2011),<br>(Avidan et al., 2014) | M | 32 | 55.3 | -2.18 | 58 | -0.21 |
| | CP<br>Mean ± s.d | | 40.4±<br>15.03 | 48.52 ±<br>18.72 | | 39.3<br>±<br>9.41 | |
| | Control<br>Mean ± s.d | | 39.3<br>±<br>13.4 | 91.57±6.24 | | 58.28<br>±<br>5.87 | |

The table shows the age and gender of participants and their performance (raw values and $z$ -normalized scores relative to a large control group) on the famous face questionnaire and CFMT. Note that 7 of the 10 CPs have participated in previous behavioral (3 CPs), and imaging (7 CPs) studies; additional behavioral measures for the CP individuals can be found in these references. Specific details regarding diagnostic and inclusion criteria can be found in the Materials and Methods section and in the related studies. The average performance on the famous faces questionnaire and the CFMT of the controls who participated in the present study is also provided (t-test comparing performance across the CP and the controls, for famous faces questionnaire $p < 0.0005$ and CFMT $p < 0.0005$ ).

**Table 5.** *Rank of nodes obtained from the comparison controls vs mild CP sub-group:*

| **a. Controls>Mild** |  | **b. Mild>Controls** |  |
| --- | --- | --- | --- |
| *Rank* | *Region Name* | *Rank* | *Region Name* |
| 1,2,3 | Anterior temporal Cortex (faces) | 1,2,3,4,5,6,9 | Right lateral occipital cortex (non-faces) |
| 4 | Right FFA (faces) | 7 | Right Superior Temporal Sulcus (faces) |
| 5 | Right Fusiform gyrus (non-faces) | 8 | Right lateral occipital cortex (faces) |
| 6,8 | Right inferior frontal gyrus opercular part (faces) | 10 | Left Superior Temporal Sulcus (faces) |
| 7 | Left TOS (non-faces) |  |  |
| 9 | Superior parietal lobule (non-faces) |  |  |
| 10 | Left Fusiform gyrus (non-faces) |  |  |

The 10 nodes with the highest node degree obtained from the controls>mild CPs (a.) and mild CPs> controls (b.) difference networks. Anatomical locations, which are based on "the atlas of human brain" (Mai et al., 2008) and validated by an expert are provided for each node.

**Table 6.** *Rank of nodes obtained from the comparison controls vs severe CP sub-group:*

| **a. Controls>Severe** |  | **b. Severe >Controls** |  |
| --- | --- | --- | --- |
| *Rank* | *Region Name* | *Rank* | *Region Name* |
| 1,2,3 | Anterior temporal Cortex (faces) | 1 | Left lateral occipital cortex (non-faces) |
| 4 | Left inferior frontal gyrus opercular part (faces) | 2,4,5,6,8 | Right inferior temporal gyrus (non-faces) |
| 5 | Left Fusiform gyrus (faces) | 3 | Left Occipital pole (non-faces) |
| 6,9 | Left Fusiform gyrus (non-faces) | 7,10 | Right lateral occipital cortex (faces) |
| 7 | Right Fusiform gyrus (non-faces) | 9 | Right lateral occipital cortex (faces) |
| 8 | Right inferior frontal gyrus opercular part (faces) |  |  |
| 10 | Right Fusiform gyrus (faces) |  |  |

The 10 nodes with the highest node degree obtained from the (a.) controls> severe (b.) severe>controls groups difference networks. Anatomical locations, which are based on "the atlas of human brain" (Mai et al., 2008) and validated by expert are provided for each node.

A direct comparison of the mild versus the severe CPs revealed similar patterns of ISFC along the anterior to posterior axis (mild > severe r_s_ = −0.38, p = 0.08, ns; Figure 4a; severe > mild r_s_ = 0.36, p=0.1, n.s., Figure 4b). Also, although not statistically significant, the patterns of results are similar to those obtained for the controls vs. the entire CPs. Specifically, the severe CPs>mild CPs contrast revealed face-specific dominance and, in the severe>mild CP group, higher ISFC patterns were associated with face-selective nodes (although it is worth noting that this is so even though the obtained network did not include the ATL). The percentage of face nodes out of the top 10 nodes (ranked in descending order by degree score) was 90% (Table 4b), compared to 35% overall (number of face nodes divided by overall number of nodes). This difference was statistically significant, χ^2^(1, n=20)=13.2, p<.001. Thus, greater severity of CP was associated with a higher number of face tagged nodes (Figure 4c). When comparing the opposite contrast (mild<severe CP), there was no statistical significance in the number of face tagged nodes, χ^2^(1, n=20)=9.89, p=n.s.

**Figure 4.**
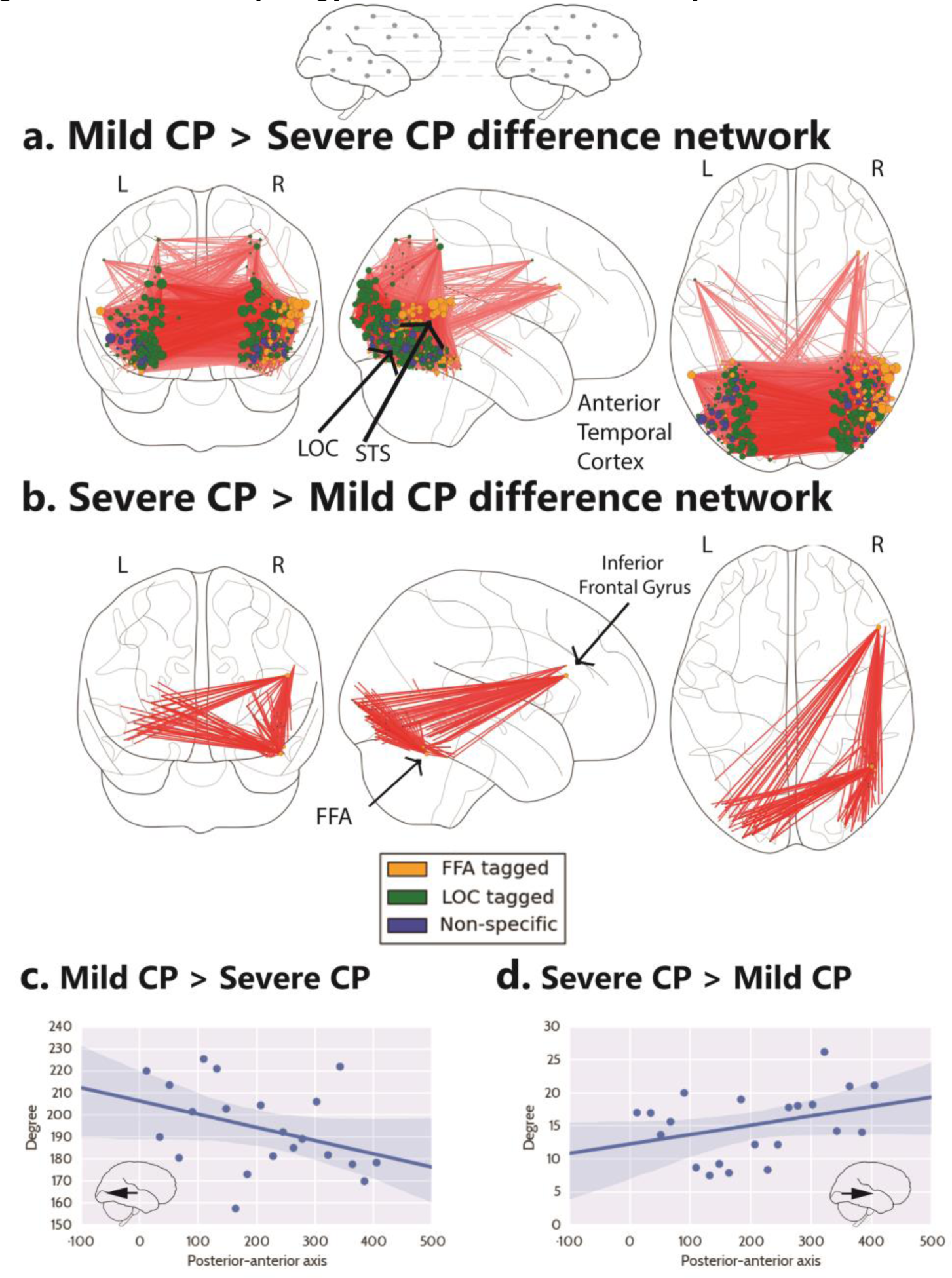
Network topology in CP is related to severity. **a.** The difference network obtained from the comparison of mild CPs > severe CPs. Conventions as in Figure 1. The mild CP group shows higher posterior ISFC compared to the severe CP group. **b.** Correlation between the degree measure of the mild CPs > severe CPs network nodes (r_s_ = −0.38, p=0.08). **c.** The difference network from severe CPs > mild CPs. As is evident, higher connectivity patterns in the severe CP group, compared to the mild CP group, are associated with face selective nodes (severe CPs >mild CPs). Importantly, however, this network does not include the ATL. d. Correlation between the degree measure of the severe CPs > mild CPs network nodes (r_s_ = 0.36, p=0.1).

**Table 4.** *Rank of nodes obtained from the comparison mild and severe CP sub-groups:*

| **a. Mild>Severe** |  | **b. Severe>Mild** |  |
| --- | --- | --- | --- |
| *Rank* | *Region Name* | *Rank* | *Region Name* |
| 1 | Left Fusiform gyrus (faces) | 1,2,8 | Right inferior frontal gyrus opercular part (faces) |
| 2,3,5 | Left Fusiform gyrus (non-faces) | 3,4,5,6 | Right inferior temporal gyrus (faces) |
| 4 | Left lateral occipital cortex (non-faces) | 7 | Right lateral occipital cortex (faces) |
| 6 | Right Middle temporal gyrus (faces) | 9 | Right Occipital Fusiform gyrus (faces) |
| 7 | Right lateral occipital cortex, inferior division (non-faces) | 10 | Left lateral occipital cortex (non-faces) |
| 8 | Right TOS (non-faces) |  |  |
| 9 | Left TOS (non-faces) |  |  |
| 10 | Left lateral occipital cortex (non-faces) |  |  |

The 10 nodes with the highest node degree obtained from the mild>severe (a.) and severe>mild (b.) CP sub-groups difference networks. Anatomical locations, which are based on "the atlas of human brain" (Mai et al., 2008) and validated by an expert are provided for each node.

Finally, to examine the right hemispheric dominance of the face tagged nodes, we applied a Mann-Whitney test on the node degree measure for each hemisphere obtained from contrasting the two groups and found a significant difference both for the severe>mild CPs contrast (Median = 10, Median = 5 in right vs. left hemisphere U = 1333, p <0.05, r = 0.17) and the mild> severe CP contrast (Median = 180, Median =107 in right vs. left hemisphere U = 1377, p = 0.05, r = 0.16) in face tagged nodes. This finding suggests that the differences in the network organization between the two subgroups of the CP individuals are more prominent in the right compared to the left hemisphere in face tagged nodes.

In sum, a direct comparison of the two CP subgroups revealed right hemispheric dominance of face-selective nodes in both cases but the two groups showed different topologies. Specifically, the mild CP group exhibited a trend towards a non-selective higher posterior ISFC pattern than the severe group and the severe CP group exhibited stronger face-related ISFC than the mild group. Importantly, in both contrasts, the obtained networks did not include the ATL.

### Estimation of the Signal to Noise Ratio of nodes

To rule out the possibility that any differences in signal to noise ratio (SNR) across groups might account for differences in network topology, a SNR analysis was conducted on the mean preprocessed raw time series of the two localizer runs in each node separately for each group. The mean of the time series was divided by the raw time series standard deviation, such that each node was characterized by a single SNR value. For each node, an independent sample t-test was conducted to compare the differences between the SNR values of the CP and control groups while correcting for multiple comparisons using an FDR correction. Critically, no significant differences in the SNR values were obtained between CP and matched controls in any of the nodes thus excluding this potential confound.

Moreover, that we observed hyper-correlation in CP relative to control subjects in posterior occipital cortex (i.e. higher reliability of stimulus locked visual responses across subjects as measured by ISFC), rules out the possibility that any differences in SNR across groups might account for differences in network topology. Rather, this finding indicates that the stimulus-locked activity patterns are reliable in both groups, despite the alterations in the network topology across the two groups.

## Discussion

The study of individuals with CP is useful from both a basic science and a translational perspective. On the one hand, CP individuals provide a unique opportunity to examine the underlying network function of normal face processing while avoiding some of the pitfalls that arise when studying patients with frank lesions (and altered vasculature) to cortex and, on the other hand, characterizing the deficits in these individuals relative to controls elucidates the atypical profile of this neurodevelopmental disorder. Also, because face recognition appears to be accomplished via the coordinated activity of multiple nodes of a distributed neural network (Haxby et al., 2000; Fairhall and Ishai, 2007; Davies-Thompson and Andrews, 2012; Gschwind et al., 2012; Joseph et al., 2012; Phillips et al., 2012; Pyles et al., 2013; Zhen et al., 2013), such a study provides the opportunity to determine whether a disrupted function of one or more of the nodes within this network leads to an altered organization of the network and, if so, to document the nature and topology of this altered organization.

By isolating stimulus-locked neural responses using a novel ISFC analysis, the present study revealed large-scale alterations in the topological structure of the visual network in CP vs control subjects.

We exploited an ISFC approach, which filters out intrinsic neural dynamics and non-neuronal artifacts that can influence FC patterns within a brain but are uncorrelated across brains of different participants (Hasson et al., 2009; Simony et al., 2016). We aimed to characterize and contrast the normally functioning face network in healthy individuals with the face network of individuals with CP who are markedly impaired at face processing. The ISFC analysis revealed a unique pattern of anterior to posterior differences in the significance of nodes in the network of CPs vs. controls. Specifically, we documented hyper-connectivity in posterior visual areas in CPs vs. controls, and hypoconnectivity between the occipital areas and anterior temporal and frontal areas. Such alterations in the topology of the correlation patterns held for both the more severely and the more mildly affected CP subgroups. Interestingly, the milder subgroup evinced a higher number of edges than the severe CP group in the posterior aspect of the network, a point we take up for further discussion below. In additional, complementary analysis revealed face-specific dominance, reflected as higher degree nodes that are face selective (defined by a localizer) in the control group compared to the CP group. In fact, in the comparison of CPs>controls, six of the top ten nodes belonged to the right inferior temporal gyrus and three nodes belonged to the lateral occipital cortex (LOC), of which only a single node was face specific. Finally, face selective nodes exhibited greater ISFC in the right versus the left hemisphere in the network obtained from contrasting CPs and controls. This pattern may be related to the well-documented hemispheric dominance of face processing. Thus, those face selective nodes in the right hemisphere, which are known to be critical for normal face processing also exhibit greater deviation from normal connectivity when contrasting CP and controls.

### The anterior temporal cortex

Perhaps the most striking difference in the topology of the face network in the controls versus CPs concerns the differences in the connectivity to the anterior temporal lobe (ATL). The ATL is implicated in the integration of person-specific information (Kriegeskorte et al., 2007b; Simmons et al., 2009; Nestor et al., 2011; Yang et al., 2014a) and familiar people recognition (Gainotti et al., 2003; Gainotti, 2007). Specifically, damage to the left ATL is more associated with impaired representation of semantic information, while damage to the right ATL impedes the visual recognition of familiar faces (Gainotti, 2007). Based on both functional activation data, functional connectivity data (during task and at rest) and structural volumetric and connectivity findings, we have argued previously that an abnormality in the anterior temporal lobe and its connectivity may play a critical role in the neural basis of CP (Behrmann et al., 2007; Avidan and Behrmann, 2009; Avidan et al., 2014). The abnormality of the ATL is evident in the current findings as well, and the analysis, conducted in an assumption-free fashion revealed that the ATL was the most important hub that distinguished the network topology of the CPs and controls. Together, these findings point to abnormal structure, function and connectivity of this region in CP individuals (but see Gomez et al., 2015; Song et al., 2015; Lohse et al., 2016).

### Altered organization of the face network in CPs

Our findings indicate that the impaired ISFC of the ATL may result in hyper-ISFC of the face-selective nodes especially the more posterior nodes, as in the right inferior temporal gyrus and the LOC, in CP compared to controls, and to a greater degree in the right versus left hemisphere. These alterations in topology offer a possible account for the face recognition deficits exhibited by CP individuals (Avidan et al., 2011; Tanzer et al., 2013; Yang et al., 2014b). For example, the posterior hyper-connectivity in CP, which is more evident in the right than left hemisphere in tandem with the residual, weak, connectivity with anterior regions may allow for some structural encoding of face stimuli derived from the immediate visual input (see Behrmann et al., 2005 for relevant behavioral findings). Although insufficient for facial recognition and individuation of identity, this rudimentary processing may partially compensate for the failure to utilize the ATL and its connectivity (Yang et al., 2014a).

Furthermore, in the current study, CP subjects exhibited greater LOC-related ISFC compared to normal subjects. Consistent with this finding, it has been demonstrated that individuals with autism spectrum disorder (ASD) who have selective deficits in face recognition show greater activation of the LOC (Schultz et al., 2000; Hubl et al., 2003). Numerous studies have shown that the LOC is associated primarily with object perception (Malach et al., 1995; Grill-Spector et al., 2001; Freud et al., 2015) and plays a major role in the processing of inverted faces, perhaps through object-like and feature-based processing (Rosenthal et al., 2016; Yovel and Kanwisher, 2005; Pitcher et al., 2011; Matsuyoshi et al., 2015).

Together, these findings offer a possible account for the fact that most CP individuals, despite their severe deficit in recognizing the identity of individual faces, are typically able to detect the presence of a face in a scene. Specifically, it is possible that the network edges located in posterior visual areas (e.g. LOC) and in the right hemisphere allow for the computation of face representations that are largely driven by the immediate visual input. This information may be relied on disproportionately by the CP individuals and perhaps serves as partial compensation for the failure to represent person-selective information which, in the normal brain is supported by the anterior temporal cortex and its connectivity (Gainotti, 2007, 2013; Fox et al., 2008). The current investigation does not allow us to infer causality, and thus, it remains to be determined whether the disconnection between the anterior and the posterior regions are the cause or the effect of the altered network organization.

### Is CP a heterogeneous disorder?

A fundamental question in the characterization of CP is whether, within the CP population, we can distinguish between subgroups that may account for the variability often found across studies (McKone et al., 2007; Russell et al., 2009; Wilmer et al., 2010). In the present study, we successfully uncovered differences between the network topology of the mild and severe subgroups when these were directly contrasted. These somewhat different profiles appear to account for the severity of the disorder. Specifically, while the network of hyper-connectivity patterns in the more posterior areas of the mild CP group may serve to compensate for the lack of connectivity to the ATL, this altered organization is less pronounced in the severe group. Furthermore, it appears that the severe group utilizes the face selective nodes (but not the ATL) to a greater extent compared to the mild CP group (see Table 4b). The use of the face-selective nodes is potentially a less effective strategy for coping with the apparent lack of connectivity of the ATL and is perhaps related to the more severe behavioral profile. The current results in which network topology and behavior are associated, are correlational in nature, and, clearly, additional studies to determine casual relations between network topology and behavior are sorely needed.

A final, related unresolved issue is whether the impairment in CP represents the lower end of the continuum of the normal distribution of face processing abilities or whether it is a separate phenomenon and, hence, a distinct disorder. The findings of a qualitatively different network in CPs compared to controls (rather than just a quantitatively different network) suggests that CP may indeed be a separate phenomenon. Further imaging studies are warranted in order to determine this critical issue.

### Conclusions

To the best of our knowledge, these data provide the first demonstration of wide-scale network topological differences in individuals with CP. Utilizing the ISFC approach, the results validated previous atypical ATL connectivity findings in CP and enabled us to gain further insight into the altered network-wise configuration of individuals with CP. We propose that investigations of the topology of other neurodevelopmental disorders might benefit from the analytic approach developed here and that insights into their underlying neural mechanisms might also be gained.

## Materials and Methods

A requirement for network analysis is the initial formulation of components of the network as a set of nodes and edges, and there are many ways in which this can be accomplished (Bullmore and Sporns, 2009; Rubinov and Sporns, 2010). Below we detail the specific approach we have taken:

### Definition of the network edges

A standard measure which is often used in fMRI studies for characterizing the edges in a functional network is the correlation coefficient between pairs of nodes identified within each subject (Smith et al., 2011). This approach was applied in the present study as well. However, despite its popularity, a major limitation of this measure, which captures within-subject synchronous activity, often referred to as functional connectivity, is that spontaneous intrinsic neuronal activity cannot be reliably separated from the evoked activity associated with the task (Greicius et al., 2003; Fox and Raichle, 2007; Simony et al., 2016). Additionally, non-neuronal artifacts such as respiratory rate and motion might also affect the results, usually by decreasing the signal to noise ratio (SNR). Indeed, in the preset study we were not able to demonstrate some of the results following correction for multiple comparison due to the limited SNR of the approach.

A possible partial solution to these limitations, formulated to increase the network inference SNR (Simony et al., 2016), relies on network construction using inter-subject functional correlation (ISFC) rather than exploiting the more widely used within-subject functional correlations. Briefly, for each pair of preselected regions of interest (ROIs) or nodes, correlation coefficients are computed between each participant and the mean signal of the remaining group without this particular participant (leave-one-out), and then correlations are averaged across all pairwise iterations. As the correlations are calculated between different participants, any intrinsic activity and artifacts, which are not correlated between participants, are cancelled out and, consequently, only the task-induced activity of interest serves as the basis of the correlations, resulting in an improved SNR.

The present study is conducted on individuals with CP, a relatively uncommon disorder and, hence, sample size is inherently limited, especially when contrasting the group size against the large number of nodes and edges (i.e., the number of variables outweighs the number of samples). This situation is typical in investigations of special populations and so we used a version of the inter-subject functional correlation (ISFC) (Hasson et al., 2009; Simony et al., 2016) modified to meet the constraints of the present scenario.

### ISFC Formulation

The current variant of ISFC, designed to take a small sample size into account, is composed of the following steps: First, each experimental group (e.g. controls and CPs) is randomly split into 2 halves and the raw signal for each node is averaged across all participants in each split. Second, for each experimental group, correlation coefficients are calculated between each pair of independent nodes, such that each node is drawn from a complementary split. Third, the correlation coefficients are transformed by Fisher z-transformation. Fourth, the groups’ positive correlation coefficients are compared. Fifth, this process is iterated N times using a bootstrap analysis to yield the null distribution and hence serves as a benchmark for deviation (where N is a large number; in the current study N is 1000). Finally, an empirical significant difference is calculated between the groups as the number of times that each correlation coefficient was larger for each group for each pair of nodes divided by the number of bootstrap iterations, while correcting for multiple comparisons using the false discovery rate [qFDR<0.005; (Benjamini and Hochberg, 1995)] procedure (see ISFC workflow in Figure 5).

**Figure 5.**
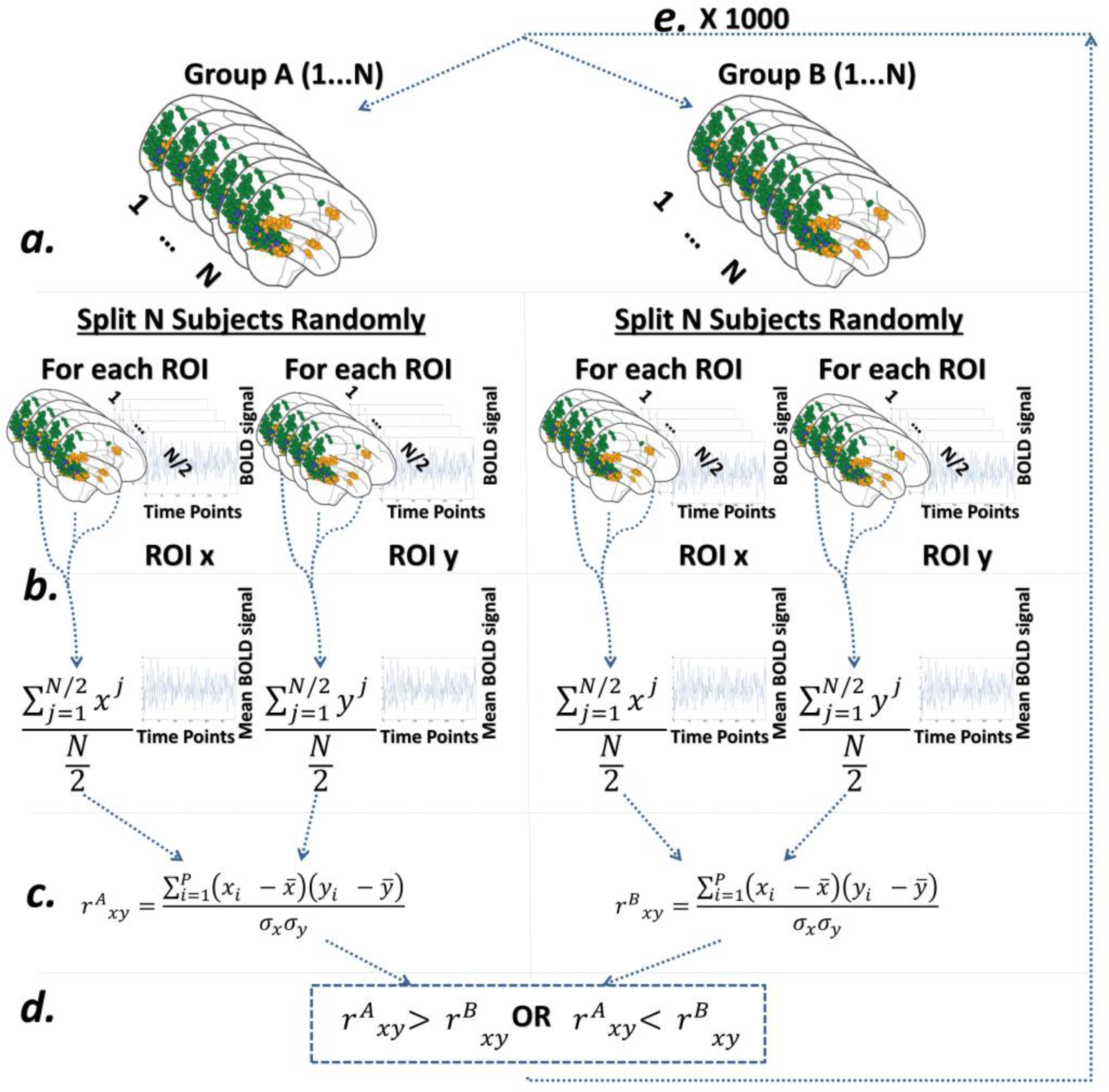
ISFC Analysis workflow. **a.** Each group of individuals (CP and controls) is randomly split into 2 halves **b.** The raw signal for each node is averaged in each split **c.** Correlation coefficients are calculated for each pair of nodes, such that each pair of nodes consists of one node from every split (the correlation coefficients are transformed by Fisher z-transformation). **d.** The correlation coefficients are compared across the groups **e**. This process is iterated N times using a bootstrap analysis (where N is a large number, in the current study 1000). Finally, an empirical significant difference is calculated between the groups as the number of times each correlation coefficient was larger for each group for each pair of nodes divided by the number of bootstrap iterations, while correcting for FDR (nodes that are group A>group B and vice versa).

### A priori localizer for node definition

#### Participants

Sixteen healthy, right-handed individuals (8 females) with normal or corrected-to-normal vision participated in the experiment (mean age ± SD = 24.5 ± 1.11). The data from three additional participants were discarded due to excessive noise. The experiment was approved by the Helsinki committee of the Soroka Medical Center, Beer Sheva, Israel and all participants signed informed consent.

#### MRI setup

Participants were scanned in a 3T Philips Ingenia scanner equipped with a standard head coil, located at the Soroka Medical Center, Beer Sheva, Israel. fMRI BOLD contrast was acquired using the gradient-echo echo-planner imaging sequence with parallel acquisition (SENSE: factor 2.8). Specific scanning parameters were as follows: whole brain coverage 35 slices, transverse orientation, 3 mm thickness, no gap, TR=2000 ms, TE = 35 ms, flip angle=90°, FOV=256×256 and matrix size 96×96. High-resolution anatomical volumes were acquired with a T1-weighted 3D pulse sequence (1×1×1 mm3, 170 slices).

#### Visual stimulation

Stimuli were presented using the E-prime 2.0 software (Psychology Software Tools, Inc., Pittsburgh, PA, USA) and projected onto an LCD screen located in the back of the scanner bore behind the subject's head. Participants viewed the stimuli through a tilted mirror mounted above their eyes on the head coil.

#### Localizer scan

A standard blocked-design localizer experiment was used to define face and non-face selective regions. Stimuli were presented in 10 sec blocks of famous faces, unfamiliar faces, buildings, daily objects, or scrambled objects (1 image was presented twice as part of the task) interleaved by 6s rest periods. Within each block there were 9 images. Each image was presented for 800 ms followed by 200 ms inter-stimulus interval and participants performed a two forced alternative choice task (see detailed description of the protocol in (Avidan et al., 2014)). The data from these participants were used to identify the nodes or regions to be used in the analysis of the CP individuals.

### Main experimental scans

#### Participants

All participants had normal or corrected-to-normal vision. The experiment was approved by the Institutional Review Boards of Carnegie Mellon University and the University of Pittsburgh, and all participants provided informed consent.

#### Congenital Prosopagnosia

Ten healthy [8 right-handed as confirmed by the Edinburgh Handedness inventory (Oldfield, 1971)] individuals diagnosed with CP (8 females, 2 males), aged between 18 and 62 years, participated in this study (mean age ± SD = 40.04 ± 15.03). None of the CP individuals had any discernible lesion on conventional MRI scanning, and none had a history of any neurological or psychiatric disease by self-report. All CP participants reported substantial lifelong difficulties with face processing. The data for 7 of the 10 CPs have been reported previously (see detailed description in Table 1). Detailed behavioral profiles were obtained and only those participants whose performance fell below 2 standard deviations of the mean of the control group on at least 2 of the 4 diagnostic measures were included [Cambridge Face Memory Test (CFMT), Famous face questionnaire, Cambridge Face Perception Test (CFPT), and a task measuring discrimination of novel upright and inverted faces; see description of the behavioral tests and details regarding prior publications for each subject in Table 1].

#### Matched controls

Ten healthy individuals, aged 25–62 years (mean age ± SD = 39.3 ± 13.4), who did not report having any difficulties with face processing participated in the imaging experiment. The CP and age-matched controls did not differ in age (p = 0.84). There was a significant difference between the control subjects and the CP group on their performance in the famous faces questionnaire (t(15) = 6.09, p<0.0005) and CFMT (t(15)= 5.54, p <0.0005; see Table 1 for mean performance), confirming the behavioral deficit in the CPs included in this study.

## Imaging Experiment

### Visual Stimulation

Visual stimuli were generated using the E-prime IFIS software (Psychology Software Tools, Inc., Pittsburgh, PA, USA) (for details see (Avidan et al., 2014)).

#### MRI Setup

Subjects were scanned either in a 3T Siemens Allegra scanner, equipped with a standard head coil (5 CPs, 4 controls) or in a 3T Siemens Verio scanner equipped with a standard head coil (5 CPs, 6 controls), using similar scanning parameters. For detailed description of the specific scanning parameters and acquisition order during the scanning session see (Avidan et al., 2014).

### Visual Stimulation Experiment

Stimuli consisted of 10 images of emotional faces (angry, fearful), neutral faces, famous faces, unfamiliar faces (Avidan and Behrmann 2008) or buildings, presented in separate 10 s blocks. Blocks were separated by 6s rest periods and there were 7 repetitions of each block type. Each image was presented for 800 ms followed by 200 ms inter-stimulus interval; participants performed a 2 alternative-forced choice task during scanning. Detailed description of the stimuli can be found in (Avidan et al., 2014).

## Data analysis

### Preprocessing of anatomical data

Anatomical scans were first preprocessed using the FSL anatomical processing script (fsl_anat) which includes the reorientation of the images to the standard (MNI) orientation (fslreorient2std), automatic cropping of the head from the image (robustfov), bias-field correction (RF/B1-inhomogeneity-correction) (FAST), registration to standard space (linear and non-linear) (FLIRT and FNIRT), brain-extraction (FNIRT-based), tissue-type segmentation (FAST) and subcortical structure segmentation (FIRST) (Jenkinson et al., 2012).

### Preprocessing of functional data

Preprocessing was conducted using dedicated Nipype pipeline (Gorgolewski et al., 2011). We utilized different components from various neuroimaging software packages including SPM8 (Penny et al., 2011), FreeSurfer (Fischl, 2012) and FSL (Jenkinson et al., 2012) as well as in-house code written in Python.

The preprocessing consisted of volume realignment of the functional scans (6 directions) to the mean EPI using SPM8 (Friston et al., 1996); artifact detection of functional scans bounded by GM mask which marked as outliers images with intensities greater than 3 standard errors from the mean intensity and images with normed composite motion differences between successive motion volumes of 1 (Gorgolewski et al., 2011); registration of functional to anatomical scans using FSL’s BBR registration procedure [epi_reg; (Greve and Fischl, 2009; Jenkinson et al., 2012)]; regressing out motion of CSF and WM first 6 components of PCA outliers [compcorr; (Behzadi et al., 2007)]; detrending (removal of 2nd order polynomials); normalization to non-linear MNI space using transformation matrices which were obtained from FSL’s anatomical preprocessing script; and, finally, spatial smoothing (6 mm) using SPM8 (Penny et al., 2011). [For related pipeline analyses used in other studies, see (Smallwood et al., 2013; Schaefer et al., 2014)].

### Definition of nodes

In order to conduct a network analysis, one needs to define a sufficiently large number of nodes that, in themselves, are relatively small. To gain a finer resolution of the network while maintaining the functional origin of each node, a seed-based correlation mask was constructed based on all voxels activated by seeds in the FFA and LOC located in the right hemisphere. These high-order visual regions were selected as seeds due to their well-documented roles in face and non-face processing respectively.

A SPM8 random-effects group analysis was conducted, such that in a first-level analysis, contrasts between stimulus categories were calculated and these were then used in a second-level analysis with subject as a random factor. The analysis included high-pass filtering with a cutoff of 1/128 Hz and a first-order autocorrelation correction. Right FFA and right LOC seeds were initially marked based on the contrasts of faces>(houses and objects) and (houses and objects)>faces using xjview, an SPM add-on tool for defining ROIs, after FDR<0.05 correction (Cui et al., 2011). The selected voxels (see below for selection procedure) were later sub-divided into small, spatially constrained clusters. This procedure preserved the original functional specificity of each voxel with "functional tagging" marking the functional preference of each node.

Using a custom Python script, voxels obtained from the functional localizer scans were limited to a gray matter mask based on the intersection of all the individual gray matter masks in MNI space (Abraham et al., 2014). Then, the time course was z-scored and a mean time course for all localizer runs averaged across all subjects was created. Using the mean time course, two separate seed-based correlation maps based on the right FFA and LOC initial seeds were defined and binarized using a 0.5 r value threshold. A union between the two maps was created and a clustering analysis was performed using the scikit-learn Ward Agglomeration algorithm (Pedregosa et al., 2011) resulting in nodes which are either face selective, non-face selective, and nodes which are equally selective for both of these stimuli (i.e. voxels which showed an overlap for both the FFA and LOC seeds at r>0.5). The upper bound of spatially constrained sub-regions was set to 500 and was determined by practical reasons with a compromise between the need for a finely grained resolution of nodes, on the one hand, and the requirement to avoid regions that are too small and might result in poor signal to noise ratio, on the other. Hence, nodes with less than 10 voxels were eliminated from the analysis, resulting in 415 nodes for the obtained networks. The total number of voxels which were removed due to this threshold of a minimum cluster size was 313 voxels out of 13407 (eliminated nodes had 3.86 voxels on average with a 2.42 standard deviation, while remaining nodes had 31.55 ± 18.4 voxels on average). Of note is that one possible outcome of this unsupervised clustering analysis is that more than a single node can reside within the “classical” functional definition (contrast based definitions) of a node such as for example, within the LOC or the FFA. This will be evident in the Results section when the characteristics of the nodes are specifically described. Visualization of networks was done with custom Python script utilizing Nilearn library (Abraham et al., 2014).

### Definition of edges

#### Standard functional connectivity (FC)

For each subject and localizer run, the time course of each of the nodes was extracted after standardization (zero mean and unit variance) and high-pass filtered using a cutoff of 1/128 Hz. Correlation matrices were constructed for each run and each subject using pairwise Pearson correlation coefficients between each pair of nodes and a Fisher r-to-z-transformation was applied to each edge. Additionally, a z normalization was applied across all correlation coefficients for each subject to remove any subject level global effects. The two localizer correlation matrices were averaged for each subject. An independent-samples t-test was conducted to compare each edge in the CP and control group while correcting for FDR (q<0.05). The outcome of this procedure are two networks which capture the significant differences in FC between Controls>CPs and the reverse, CPs>controls.

### ISFC

#### Split Group analysis based to the main experimental scans

For each subject and localizer run, the time course of each of the nodes defined in the separate localizer scan was extracted after standardization (zero mean and unit variance) and high-pass filtered using a cutoff of 1/128 Hz. The two localizer scans were averaged. Split group analysis (ISFC) was performed on the average time course while contrasting the CP and matched controls group.

### Construction of difference networks

Two difference networks which capture the significant difference in ISFC between Controls>CPs and CPs>controls were constructed. This was done for each of these comparisons separately in the following manner: Using the ISFC procedure, any edge that was not empirically significantly different between the two groups across the 1000 bootstrapped iterations, while correcting for FDR, was removed from the analysis and was set to zero. For visualization purposes, all the remaining significantly different edges were given a weight that corresponds to the mean value of the difference between the different groups (Controls>CPs and CPs>Controls) across all 1000 iterations of the ISFC (See Figure 1 and Figure 5 for ISFC workflow).

## Footnotes

1 To whom correspondence should be addressed.

## Author contributions

G.R, G.A and M.B designed and conducted the research; U.H and E.S: contributed novel analytic tools and provided insights on their adaption for the present study; G.R developed the data analysis approach and G.R, G.A and M.T analyzed data; G.R, G.A, and M.B. wrote the manuscript.

The authors declare no conflict of interest.

## Acknowledgments

This work was support by ISF grant 296/15 to GA and NSF grants BCS-1354350 and #SBE-0542013 to MB

